# Functional Templates in fMRI: Building Accurate and Interpretable Group-Level Decoders

**DOI:** 10.64898/2026.05.21.726781

**Authors:** Pierre-Louis Barbarant, Florent Meyniel, Bertrand Thirion

## Abstract

Inter-individual variability poses a significant challenge in decoding brain activity across subjects. While standard anatomical registration procedures reduce morphological differences, they fail to capture functional variability between subjects. Functional alignment methods address this issue by establishing voxel-to-voxel correspondences between pairs of individuals, thereby constructing a shared functional space. This shared space can be extended at the group level by generating a functional template. However, despite the availability of toolboxes, functional templates remain underused in fMRI analysis. Adopting this approach is currently difficult due to the diversity of existing methods and the lack of clear guidelines. Comprehensive evaluations of functional templates are limited to movie-watching paradigms. Here, we extensively compare functional alignment methods (Optimal Transport, Procrustes, Ridge regression, and Shared Response Model) and template construction strategies (in-sample, out-of-sample, pairwise) within the more general framework of task decoding. In this framework, decoding accuracy measures how well individual activation patterns align. Across multiple tasks and datasets, we demonstrate that population templates built using Optimal Transport (a) yield the highest decoding accuracy, (b) are not significantly biased by individual subjects, which facilitates generalization to new subjects, and (c) preserve the cortical signal topography.

## 1 Introduction

In functional magnetic resonance imaging (fMRI), comparing activity patterns across individuals presents a fundamental challenge due to variations in both brain anatomy and function. This challenge is particularly acute because fMRI captures both structural differences in brain organization and individual variations in functional responses. While traditional anatomical normalization approaches address this by mapping individual brains onto standard templates through various transformations (Fonov et al., 2009), these methods do not adequately capture the full scope of functional differences between subjects (Haxby et al., 2001; Sabuncu et al., 2009).

### 1.1 Functional Alignment: Pairwise vs. Template Strategies

Functional alignment has emerged as a complementary approach to anatomical registration and spatial normalization. It offers methods for establishing correspondences between neural patterns across individuals using fMRI maps. Current techniques generally fall into two categories: pairwise and template-based.

Pairwise methods, including Procrustes Analysis (Guntupalli et al., 2016) and Optimal Transport (OT)-based alignment (Bazeille et al., 2019), directly map functional responses between pairs of subjects. While effective, these methods can only decode a given target subject; therefore, they lack a unified space for group analysis. Furthermore, these methods encounter computational challenges as the number of subjects increases.

Template-based approaches address these limitations by creating a shared representational space between all subjects. Methods such as Generalized Procrustes Analysis (Gower, 1975) and Hyperalignment (Guntupalli et al., 2016) iteratively refine a common functional template across all subjects. This eliminates the need for an arbitrary reference subject and has shown promise in improving group-level analyses (Jeganathan et al., 2024).

However, template-based functional alignment introduces two important challenges. First, jointly estimating the individual transformations and the group template is computationally demanding, as each refinement of the template requires recomputing every subject-to-template transformation. Second, evaluating how well a template generalizes to new subjects, as emphasized in Guntupalli et al., 2016, requires a leave-one-subject-out cross-validation scheme, which leads to a quadratic number of transformations. Together, these costs limit the feasibility of the approach for studies with limited computational resources.

The Shared Response Model (SRM) (Chen et al., 2015) represents another significant advancement, with implementations like FastSRM (Richard & Thirion, 2023) that offer computational efficiency and improved decoding performance (Bazeille et al., 2021). However, SRM’s reliance on latent space representation limits interpretability. The template has no representation in voxel space, which prevents activity mapping on the cortex. In contrast, functional templates in image space maintain direct correspondence with brain anatomy, providing interpretable composite representations of neural patterns across subjects. Given their anatomical interpretability, image-space templates are particularly valuable for neuroscientific investigations, as they can be analyzed using traditional neuroimaging approaches and compared directly with anatomical templates.

### 1.2 Key Open Questions

Despite the expected benefits of functional templates, several critical questions remain unexplored. First, there has been limited systematic comparison between template-based and pairwise alignment strategies for a popular analytic approach in functional brain imaging: decoding conditions or variables of a task (hereafter “task decoding”). In particular, voxelwise inter-subject correlation is the most reported metric in previous studies comparing different alignment methods (Andreella & Finos, 2022; Guntupalli et al., 2016, 2018; Haxby et al., 2020), yet it is highly sensitive to spatial blurring and poorly informative in task settings (Bazeille et al., 2019; Chen et al., 2015). It remains unclear whether template-based methods offer clear advantages over pairwise alignment in the case of task decoding. Second, the extent to which voxel-space functional alignment preserves brain topography–for example, the precise localization of relevant voxels on the cortex–relative to anatomical registration requires investigation. Third, including test subjects in template generation could bias generalization, as noted by Jeganathan et al., 2024 for Procrustes; such a bias needs to be evaluated across different alignment methods.

### 1.3 Our Contributions

This study addresses these knowledge gaps through a comprehensive evaluation framework with three key contributions:

1. **Systematic comparison of alignment strategies** We evaluate three distinct strategies: out-of-sample template (the test subject is excluded from template generation, Chen et al., 2015; Jeganathan et al., 2024), in-sample template, and pairwise alignment across multiple alignment methods (Optimal Transport, Procrustes, Ridge regression, and Shared Response Model). Unlike previous benchmarks that relied on movie-watching data (Chen et al., 2015; Guntupalli et al., 2018; Jeganathan et al., 2024) or separate task data for alignment and decoding (Bazeille et al., 2021), we use the same task data for both alignment and decoding (with cross-validation). This design makes functional alignment more practical by eliminating the need for external data to compute the alignment, and it isolates inter-subject variability in the data relevant for decoding from other factors (present in external data).
2. **Empirical assessment of template bias**: We test whether including test subjects during template generation artificially inflates decoding accuracy, comparing in-sample versus out-of-sample template strategies across all methods. These two strategies have different implications. In-sample template estimation is easier to compute and produces a single, unified template. However, it risks incorporating information from the test subject, which can artificially boost decoding performance. In contrast, out-of-sample estimation avoids this bias by ensuring templates are agnostic to the test subject. However, this method requires substantial computation due to repeated template estimation.
3. **Analysis of spatial specificity**: We examine how different in-sample alignment methods affect the spatial distribution of classifier weights in image space, revealing distinct patterns of how each method influences brain topography. This analysis demonstrates that template-based approaches enhance spatial specificity while maintaining interpretability.

Our empirical results establish clear practical guidelines: in-sample Optimal Transport templates offer unbiased alignment with superior decoding performance and interpretability. We validate these findings across a diverse range of tasks from three datasets, encompassing various cognitive domains and sample sizes.

### 1.4 Related Work

*Hyperalignment*, introduced by Haxby et al., 2011, pioneered functional alignment by constructing a shared high-dimensional representational space across subjects. Unlike generalized Procrustes analysis (GPA) (Gower, 1975), which jointly estimates all transformations in an order-invariant manner, Hyperalignment proceeds in two sequential steps and is therefore sensitive to the order in which subjects are processed. First, the method iteratively aligns each subject to a growing functional template, updating the template at each step. Second, once the template is finalized, subject-specific transformations are recomputed with respect to this reference.

The approach is typically implemented within a searchlight framework (Kriegeskorte et al., 2006), where Procrustes transformations are estimated locally on small cortical patches, and these local solutions are then aggregated into whole-brain transformation matrices. Thanks to the relatively low computational cost of Procrustes alignment, Hyperalignment is often applied in an out-of-sample manner using a leave-one-subject-out strategy. In our benchmark, we rely on GPA rather than Hyperalignment to avoid the order effect inherent to Hyperalignment’s sequential procedure, as in Bazeille et al., 2021 and Jeganathan et al., 2024.

*Connectivity-based Hyperalignment* (Guntupalli et al., 2018) extends this framework by incorporating functional connectivity information, leading to hybrid hyperalignment approaches (Busch et al., 2021) that leverage both local activation patterns and connectivity profiles. This approach is highly relevant when used in conjunction with BOLD time series data, but is outside the scope of our study as we target homogeneous sources of data, i.e, we only incorporate z-scored condition maps.

*The ProMises* model (Andreella & Finos, 2022) recasts the Procrustes alignment problem within a statistical framework. By imposing a whole-brain anatomical prior, it overcomes the locality constraints inherent to searchlight (Guntupalli et al., 2016) and piecewise (Bazeille et al., 2021) hyperalignment, while retaining a relatively low computational cost. Although comparing our framework with ProMises would be valuable, analogous statistical reformulations do not currently exist for Optimal Transport or Ridge alignment, preventing a meaningful comparison across all methods.

*Integrated Alignment* (Jeganathan et al., 2024) proposes that combining anatomical and functional alignment yields improved decoding performance, independent of the specific alignment method. The approach interpolates between the identity transformation on anatomically registered data and the functional alignment solution, producing templates that jointly capture shared structure and subject-specific functional features. However, this interpolation depends on a hyperparameter that must be tuned based on an external cohort, and the method is built within the surface-based diffeomorphic registration framework *MSMSulc* (Glasser et al., 2016), making it impractical to apply directly in our setting.

*FUGW* (Fused Unbalanced Gromov–Wasserstein) (Thual et al., 2022) formulates inter-subject alignment as a quadratic Optimal Transport problem that jointly optimizes functional similarity and spatial constraints. Beyond the functional matching term similar to the formulation of Bazeille et al., 2019, FUGW incorporates a Gromov–Wasserstein objective that aligns the pairwise geodesic structure of each subject’s anatomy. This quadratic matching of intra-subject distances enables the model to capture complex spatial correspondences that cannot be expressed by simple pointwise or linear constraints. However, solving the Gromov–Wasserstein problem is computationally demanding and scales poorly to whole-brain volumetric data. In addition, the unbalanced formulation allows for partial mappings between subjects, but introduces several extra hyperparameters. As a result, the method requires tuning three coupled regularization terms, making it difficult to deploy in the real-world large-scale scenarios considered in this benchmark.

## 2 Materials and Methods

In this section, we detail the components of our benchmark. We begin by formally defining the alignment methods used to map data between pairs of subjects (Sections 2.1 & 2.2), followed by our piecewise strategy, which ensures computational tractability (Section 2.3). We then describe the group-level strategies for constructing functional templates (Section 2.4) and present the three datasets on which our evaluations are conducted (Section 2.5). Finally, we outline the design and motivation of the experimental setup used throughout the benchmark (Sections 2.6 & 2.7).

### 2.1 Notations

Inter-subject functional alignment seeks to establish voxel-wise correspondences across individual brains to improve group-level analyses. Although we operate in the volumetric setting, the mathematical framework is agnostic to the volume–surface distinction; all methods considered here can be equivalently applied on cortical surface meshes by treating vertices in place of voxels, as in Jeganathan et al., 2024. Throughout, we assume that all subjects have already been anatomically normalized to a common template space.

Let *p* denote the number of voxels (or vertices) shared across subjects after anatomical normalization, and let *n* denote the number of task-related effect maps (hereafter “task maps”) used for alignment, also consistent across subjects. For each individual, we stack the *n* task maps into a feature matrix **F** ∈ R*^n^*^×^*^p^*. All matrices maintain a similar ordering between tasks and voxels respectively across all subjects. We will denote by ||.||*_F_* the Frobenius norm over matrices.

### 2.2 Pairwise Alignment Methods

In this section, we will consider the two-subjects alignment case. We denote one subject as the *source* with a corresponding feature matrix **F***^s^* and the other as the target **F***^t^*. We seek out to compute a functional mapping from the source to the target subject.

#### 2.2.1 Procrustes Analysis

Originally developed within the *Hyperalignment* framework (Haxby et al., 2011), Procrustes analysis (Hurley & Cattell, 1962) computes an orthogonal transformation matrix **M** and a scaling factor *s* that optimally align source and target task maps. This approach treats functional patterns as rigid objects, ensuring that the transformed source data preserves distances while minimizing discrepancy with the target subject. The optimization problem is:

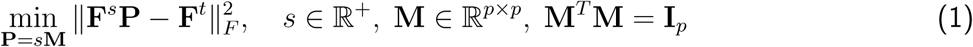

where **I***_p_*is the *p*-dimensional identity matrix, ensuring orthogonality of **M**.

#### 2.2.2 Optimal Transport

Optimal Transport (OT) was first introduced for inter-subject alignment in neuroimaging by Bazeille et al., 2019. Unlike Procrustes analysis, which assumes a scaling of the data and a rigid transformation (e.g the combination of a translation and a rotation or a reflection), OT computes a coupling a.k.a. transport plan **P** ∈ R*^p^*^×^*^p^* that minimizes the cost of mapping voxel-wise functional responses from source to target subject. A coupling is *balanced*, i.e. it ensures that each voxel from the source and target space has equal importance in the mapping. The cost matrix **C** ∈ R*^p^*^×^*^p^* encodes the dissimilarity between voxel activations of both subjects by measuring the Euclidean distance between activation vectors over every pair of source and target voxels. An optimized transport plan estimated without entropic regularization converges to a permutation matrix, indicating that non-regularized OT produces orthogonal solutions similarly to Procrustes alignment with additional sparsity constraints. Additionally, an entropic regularization term Cuturi, 2013 ensures computational tractability and smoother mappings. The optimization problem is formulated as such:

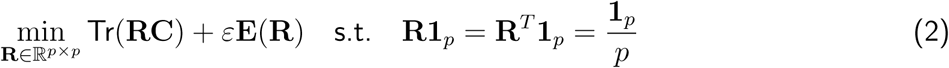

where **E**(**R**) represents the entropic penalty:

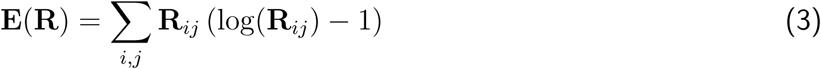

and *i, j* index source and target subjects, respectively. We fix the parameter *ε* to 0.1 to match the settings of Bazeille et al., 2021. Given the Optimal Transport plan **R**^∗^, the alignment matrix is defined by 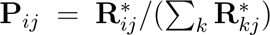As in Bazeille et al., 2019, every voxel supports the same weight, thus 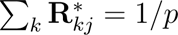 and the previous expression simplifies into **P** = *p***R**^∗^.

#### 2.2.3 Ridge Regression

Ubiquitous in fMRI encoding models (Naselaris et al., 2011), Ridge regression has recently been successfully applied to inter-subject alignment (Ferrante et al., 2024). This method imposes the mildest constraints, applying only *L*_2_ regularization to the correspondence matrix:

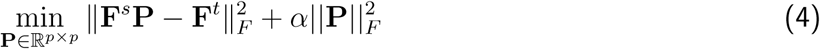

The hyperparameter *α* is tuned via cross-validation using scikit-learn’s RidgeCV implementation. Notably, as pointed out by Barbosa et al., 2025, Ridge regression differs from both Procrustes and OT in that it does not define a metric between subject activations and is not symmetric.

### 2.3 Piecewise Alignment Approaches

Aligning whole-brain voxel data across subjects poses significant computational challenges due to the high dimensionality of neuroimaging data. Furthermore, simultaneous alignment of all voxels across the entire brain may produce erroneous correspondences between anatomically distant regions. To address these issues, we adopt a piecewise alignment approach following Bazeille et al., 2021 and Jeganathan et al., 2024. This strategy leverages a brain atlas to define anatomically coherent parcels, within which voxel-wise transformations are computed independently.

We use the Schaefer atlas (Schaefer et al., 2017) for our main experiments with 400 parcels, resulting in an average of 84 voxels per parcel. Control analyses showed that the effect of the number of parcels on our conclusions is negligible (see Appendix Figure 10). This partitioning ensures localized functional alignment while maintaining computational feasibility. We provide an overview of the list of hyperparameters to fix for each method in Table 1.

**Table 1:** Free Parameters. We list the number of free parameters to set for each alignment method under each strategy. *N_parcels_* denotes the number of parcels to choose in the atlas, *ε* the entropic regularization parameter for OT, *α* the *L*^2^ penalty for Ridge regression and *K* the number of latent components for the Shared Response Model.

|  | Procrustes | Optimal Transport | Ridge | SRM |
| --- | --- | --- | --- | --- |
| Pairwise | $N_{parcels}$ | $N_{parcels}, \varepsilon$ | $N_{parcels}, \alpha$ | – |
| Template | $N_{parcels}$ | $N_{parcels}, \varepsilon$ | $N_{parcels}, \alpha$ | $N_{parcels}, K$ |

### 2.4 Template-Based Alignment

#### 2.4.1 Variational Formulation

To create a functional template serving as a shared reference space across subjects, we follow the approach of Jeganathan et al., 2024. The template is constructed in a piecewise manner to ensure interpretability in image space. Mathematically, the template is defined as the solution to the following variational problem:

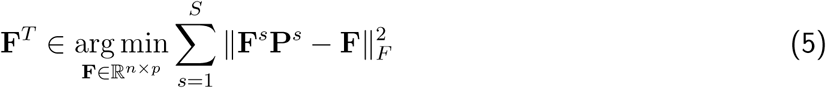

where *S* represents the number of subjects contributing to the template, **F***^s^* denotes the functional data (here, task maps) of subject *s*, and **P***^s^* is the alignment map from subject *s* to the template.

Template construction follows an alternating minimization scheme, illustrated in the left panel of Figure 1. Starting from an initial estimate, typically the Euclidean average of all subjects’ data, the procedure alternates between two steps: (i) estimating subject-specific alignment maps **P***^s^*, and (ii) projecting each subject onto the current template space and averaging the aligned data to update the template. These steps are repeated until the template converges. This general framework appears under different names depending on the alignment method, including Generalized Procrustes Analysis (GPA) Gower, 1975 for Procrustes alignment and Wasserstein barycenters Agueh and Carlier, 2011 for Optimal Transport–based alignment.

**Figure 1:**
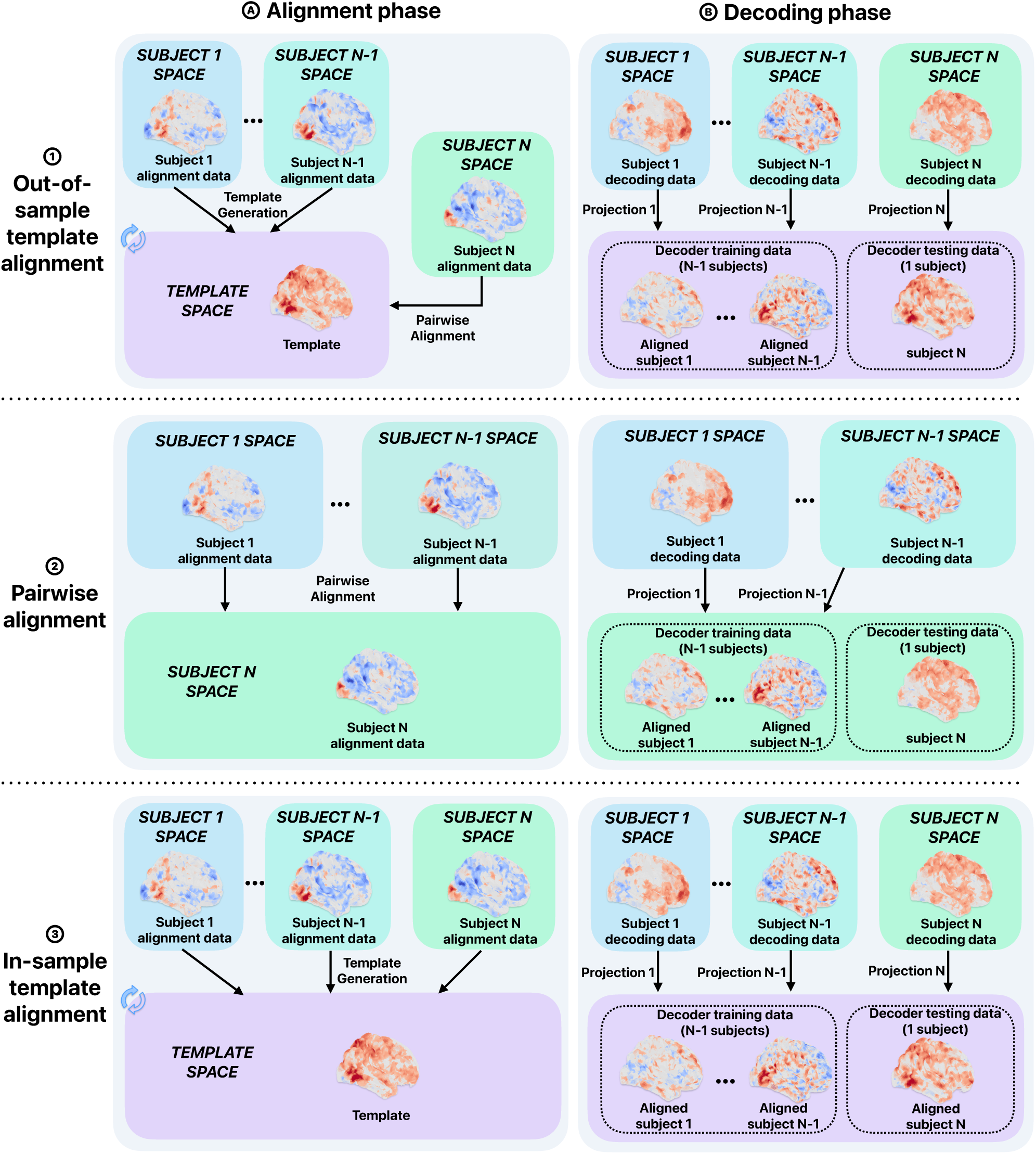
Schematic of the three alignment strategies. Each alignment strategy (out-of-sample, pairwise, in-sample) is evaluated across all combinations of tasks and alignment methods (Procrustes, Optimal Transport, Ridge regression, Shared Response Model) through a nested cross-validation procedure. Each subject’s task data is first split between alignment and decoding groups using a 5-fold outer cross-validation loop (20% alignment data, 80% decoding data). Inside each fold, the pipeline consists of two phases: **A.**an *alignment phase* where functional alignment transformations are computed from alignment data, and **B.** a *decoding phase* where these transformations are applied to separate decoding data to perform leave-one-subject-out cross-validation, yielding one accuracy score per subject. For each task and method combination, accuracy scores are averaged across the 5 folds for each subject. The three alignment strategies differ in how the functional alignment target is constructed: **1. Out-of-sample template alignment**: For each held-out test subject, a template is generated from the N-1 training subjects’ alignment data, explicitly excluding the test subject. The test subject’s decoding data is then aligned to this out-of-sample template in a pairwise fashion. The classifier is then trained on the training subjects’ aligned data and tested on the aligned test subject’s data. **2. Pairwise alignment**: For each inner cross-validation fold, the left-out test subject serves as the alignment target. All N-1 training subjects’ decoding data are individually aligned to the test subject’s space. The classifier is trained on the aligned training subjects and is evaluated on the test subject in its native space. **3. In-sample template alignment**: A single population template is computed from all subjects’ alignment data. Each subject’s decoding data is then projected to this template space using the computed transformations. Leave-one-subject-out cross-validation is performed in the common template space, where the classifier is trained on N-1 subjects and tested on the remaining subject. Technical implementation leverages fmralign’s parallel processing capabilities for template construction and alignment computations, while classification is performed using scikit-learn’s LinearSVC with default parameters.

Note that that regardless of the alignment method used, these alternate iterations are guaranteed to converge towards a result minimizing (5). However, Bazeille et al., 2019 notes that this result is not necessarily a global optimum.

#### 2.4.2 The Shared Response Model (SRM)

Thus far, we have assumed the template resides in image space. However, motivated by the search for a common lower-dimensional basis between subjects, Chen et al., 2015 proposed the *Shared Response Model* (SRM). We focus here on the deterministic approach (see Richard and Thirion, 2023 for a thorough review of modern SRM variants).

The template formulation remains as in Equation 5, except that the template now resides in a lower-dimensional space of dimension *K* ≪ *p*. Each subject-wise transformation matrix **P***^s^* ∈ R*^p^*^×^*^K^* must be adapted while preserving the orthogonality constraint (**P***^s^*)*^T^* **P***^s^* = **I***_K_*. The template is initialized from a random Gaussian distribution and computed in a piecewise manner. Given the trade-off between the number of parcels and the optimal number of latent components, we rely on the sensitivity study by Bazeille et al., 2021 and select *K* = 20 for our 400-parcel configuration.

The implementation of the aforementioned alignment methods and pairwise/template strategies is publicly available through our *fmralign* package^1^.

### 2.5 Datasets and Preprocessing

We replicate our experiments across diverse tasks from three datasets:

**Individual Brain Charting (IBC)** The IBC dataset (Pinho et al., 2024) comprises recordings from 13 participants across 80 task conditions. We selected a subset of five tasks, detailed in Table 3. These classical tasks consist of successive trials that can be labeled by their corresponding condition. We performed single-trial analysis using Nilearn (Abraham et al., 2013) to derive trial-level z-score maps, following the design of Bazeille et al., 2021. fMRI data were preprocessed using *Pypreprocess*, including distortion correction, motion correction, and spatial normalization.

**StudyForrest Music Listening** The music listening subset of the StudyForrest dataset (Hanke et al., 2015) includes 10 participants listening to audio clips from five music genres, processed following Bazeille et al., 2021.

**CNeuroMod-THINGS** The CNeuroMod-THINGS dataset (St-Laurent et al., 2025) features three subjects observing naturalistic images from the THINGS dataset (Hebart et al., 2019). We used the 27 meta-classes provided with THINGS to derive labels. However, since these meta-classes are overlapping (e.g., animal, bird, insect), we retained only four semantically distinct classes: animal, clothing, food, and vehicle. Since each image is presented only three times per subject, we derived image-level z-score maps.

For all tasks, we ensured that conditions and stimuli were matched between subjects and that classes were balanced within each task. anatomical registration was performed to the 3mm *MNI152NLin2009cAsym* volume template (Fonov et al., 2009) to ensure spatial consistency across subjects. Whole-brain masking and standardization were applied using *Nilearn* Abraham et al., 2013.

### 2.6 Experimental Design

#### 2.6.1 Task-Based Decoding Analysis

To compare the various alignment techniques, we implemented a between-subject classification of task conditions (illustrated in Figure 1, right panels). The evaluation protocol follows a leave-one-subject-out cross-validation strategy, as described in Bazeille et al., 2021, to assess decoding performance across individuals.

However, unlike Bazeille et al., 2021, which used data from different tasks for alignment and decoding, we used the same tasks for both, splitting each subject’s task data between alignment (20%) and decoding (80%), stratified by run to ensure equal contribution from each run. This choice limits sources of variability beyond inter-subject differences, notably inter-run and inter-task variability. The data split is repeated in a nested five-fold cross-validation, and results are averaged across folds.

We employ a linear support vector classifier, which has proven effective for fMRI data (Lee et al., 2010). Specifically, we use *Scikit-learn*’s LinearSVC implementation (Pedregosa et al., 2011), consistent with Bazeille et al., 2021 and Jeganathan et al., 2024.

#### 2.6.2 Three Alignment Strategies

We investigate three functional alignment strategies from the literature: two template-based approaches and one relying solely on pairwise alignments. We present them in the following paragraphs.

**Out-of-Sample Template** We adapt this strategy from Jeganathan et al., 2024 (panel 1 of Figure 1), where it was used to study template generalization to new individuals. We first set one subject apart as the left-out subject, compute a template from all other subjects, then align the left-out subject to this template in a pairwise fashion. This ensures that no information about the left-out subject is incorporated during template creation, unlike the in-sample template. However, performing complete LOSO cross-validation among *S* subjects requires generating *S* templates and performing *S* pairwise alignments, making this the most computationally intensive strategy.

**Pairwise Alignment** Used in Bazeille et al., 2021, this approach (panel 2 of Figure 1) minimizes the domain shift between subjects by mapping each subject to the left-out subject. While avoiding template creation, this strategy requires *S*(*S* − 1) mappings for complete evaluation across *S* subjects.

**In-Sample Template** Coined as “In-sample Template” by Jeganathan et al., 2024, this is the most straightforward approach (see panel 3 of Figure 1). A single template is computed from alignment data across the entire population, yielding individual mappings between each subject and the template. These mappings align decoding data to the template space, where leave-one-subject-out (LOSO) cross-validation is performed. The computational complexity in terms of alignment operations grows linearly with the number of subjects.

### 2.7 Experiments

We conducted three experiments to evaluate decoding performance across alignment methods and strategies. In all cases, alignment and decoding were performed on disjoint data splits, and results were averaged over a 5-fold, run-stratified cross-validation scheme.

**(1) Global comparison under out-of-sample template alignment.** In this first experiment, we compare each functional alignment method against an anatomical baseline (i.e., no functional alignment). For every method, template estimation is achieved using the out-of-sample scheme to avoid bias. For evaluation, classifiers are trained on the aligned training subjects projected onto the corresponding template and then tested on the held-out subject (Figure 1, row 1, panel B), yielding one decoding score per subject.
**(2) Pairwise versus out-of-sample template alignment.** We next quantify the performance gap between pairwise alignment and out-of-sample template alignment (Figure 1, rows 2 and 1, respectively). Results for the out-of-sample template condition are identical to Experiment 1. In the pairwise condition, the left-out test subject serves as the alignment target for all other subjects.
**(3) Assessing potential bias in in-sample templates.** We then evaluate whether in-sample template construction introduces bias by comparing it to the out-of-sample setting. We define the empirical *bias* of an alignment method as the difference of test scores between in-sample and out-of-sample template strategies.
**(4) Visualizing and interpreting template alignments.** Finally, inspired by Bazeille et al., 2021, we qualitatively examine how functional alignment alters the spatial distribution of decoder weights. Whereas Bazeille et al., 2021 focuses on the localization of functional signal, we instead analyze the localization of the decoder’s discriminative weights, as these directly underlie the decoding performance assessed in the previous experiments. We focus on the *face adult* condition from the FaceBody task in the IBC dataset and apply a procedure analogous to Experiment 3 to identify the voxels that most strongly drive classification of the condition. We also examine the distributions of a sparsity criterion across aligned condition maps for the various methods and compare them to anatomical alignment as a baseline.

## 3 Results

We compare the three alignment strategies in the following sections. Results are averaged across alignment/decoding data splitting folds to obtain one accuracy score per subject for each task/strategy/alignment method combination. Since performance for each method is derived from leave-one-subject-out cross-validation, subject accuracy scores are *not* independent from one another, precluding the use of standard *t*-tests to compare mean accuracies between methods. We therefore use the corrected *t*-test of Nadeau and Bengio, 1999 to assess the statistical significance of differences between methods.

### 3.1 OT-Based Templates Boost Decoding Accuracy

Figure 2 presents the results of our task classification experiment using the out-of-sample template strategy. Optimal Transport templates demonstrate a clear performance advantage, outperforming the anatomical baseline in overall accuracy on all tasks, with significant improvements in five out of seven tasks (*p* ≤ 0.05). Other methods never improved significantly over the anatomical baseline. Specifically, out-of-sample Procrustes demonstrates significantly *worse* average performance (*p* ≤ 0.05) on three datasets, Ridge regression on six datasets, while the SRM model remains capped at chance-level performance.

**Figure 2:**
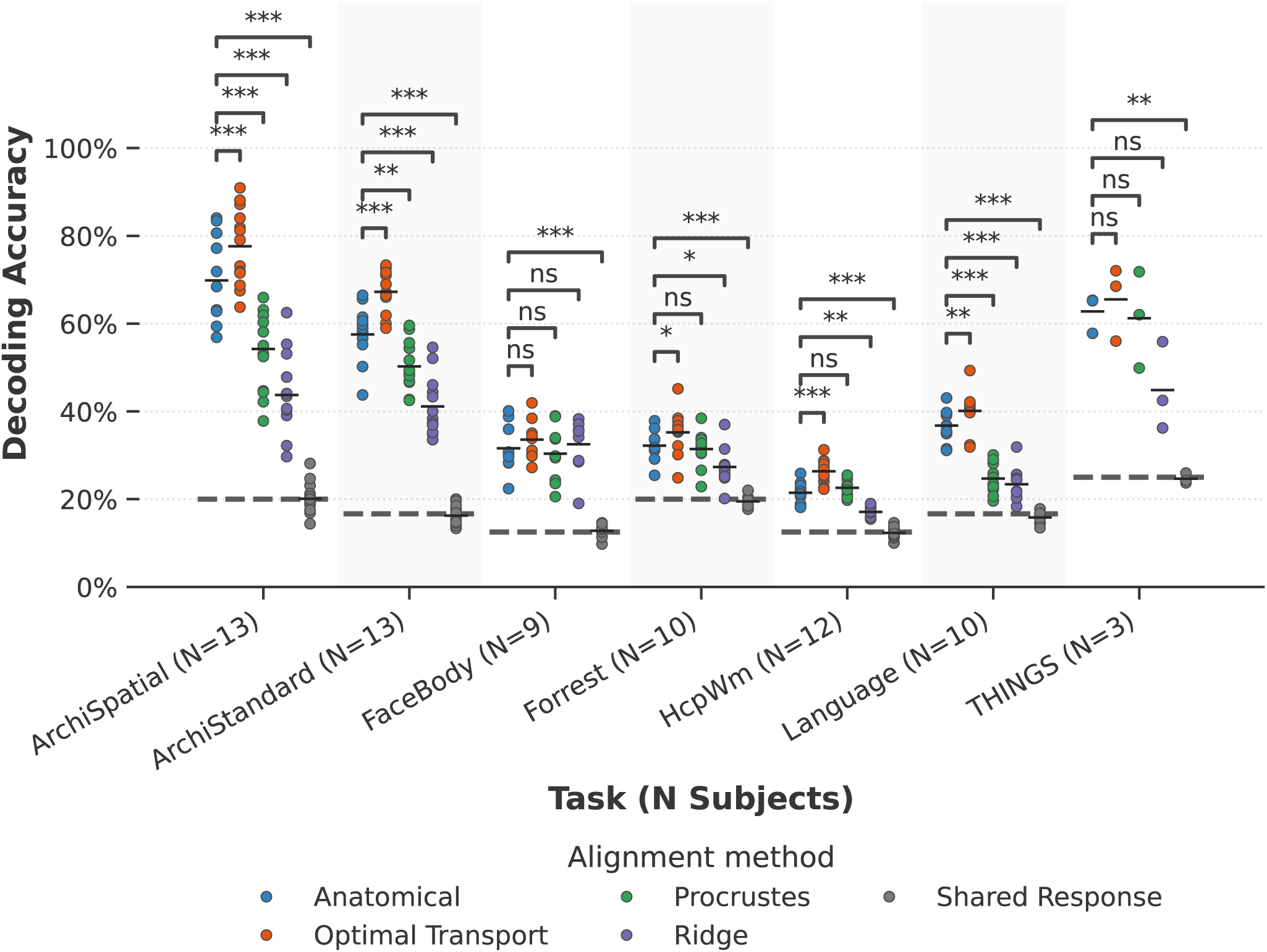
Decoding performance using the out-of-sample template. Classification accuracies are reported for each alignment method using leave-one-subject-out cross-validation. Each data point represents the decoding accuracy for a single subject, with colors indicating the alignment method. For template-based methods (Optimal Transport, Ridge, Procrustes, and Shared Response), templates are computed by leaving out each test subject (out-of-sample strategy – Panel 3 of Figure 1). Anatomical alignment serves as a baseline, using only registration to a standard anatomical template without functional alignment. Horizontal lines within each distribution represent mean accuracy values. Dashed horizontal lines indicate chance-level performance for each task. Statistical comparisons between in-sample and out-of-sample strategies are performed separately for each alignment method using *t*-tests with Nadeau and Bengio, 1999 correction. Significance levels: *\* p* ≤ 0.05, *\*\* p* ≤ 0.01, *\*\*\* p* ≤ 0.001.

Beyond mean performance, a major concern when incorporating new methods into fMRI studies is the occurrence of failure cases: instances where decoding scores fall below baseline. We report the number of subjects exhibiting decoding accuracy lower than anatomical alignment in Table 2. OT templates result in the lowest number of failures by far, with only five cases of below-baseline performance, followed by Procrustes analysis with 50 failure cases.

**Table 2:** Failure Cases. We report the number of subjects whose decoding accuracy is lower after functional alignment on the population template for each method (compared to the standard anatomical alignment). The best performance are highlighted in bold.

| Task | N Subjects | Optimal Transport | Procrustes | Ridge | Shared Response |
| --- | --- | --- | --- | --- | --- |
| ArchiSpatial | 13 | **0** | 13 | 13 | 13 |
| ArchiStandard | 13 | **0** | 12 | 13 | 13 |
| FaceBody | 9 | 3 | 5 | 5 | 9 |
| Forrest | 10 | 1 | 5 | 9 | 10 |
| HcpWm | 12 | **0** | 3 | 12 | 12 |
| Language | 10 | **0** | 10 | 10 | 10 |
| THINGS | 3 | 1 | 2 | 3 | 3 |
| Total | 70 | 5 | 50 | 65 | 70 |

These results underline the necessity of comparing alignment methods not only among themselves but also to the anatomical reference in order to assess whether they provide a genuine improvement in signal quality for across subjects. In these experiments, the reliability of OT across individuals therefore provides strong support for its use in functional alignment.

### 3.2 OT Templates Achieve the Closest Performance to Pairwise Alignment

To assess the benefits of directly aligning the decoder’s training data to the test subject’s domain, we compare the performance gap between pairwise and in-sample template approaches in Figure 3. Since the Shared Response Model is inherently template-based and thus inadequate for pairwise alignment, we exclude it from this comparison. As expected (Bazeille et al., 2021), pairwise alignment performed better than the out-of-sample template alignment. However, the performance gap was typically smaller for OT compared to other methods (see Appendix Figure 8). In particular, pairwise strategies perform best for Ridge regression, significantly outperforming (*p* ≤ 0.05) out-of-sample template decoding on all tasks, followed by Procrustes analysis on five tasks and OT on only two tasks.

**Figure 3:**
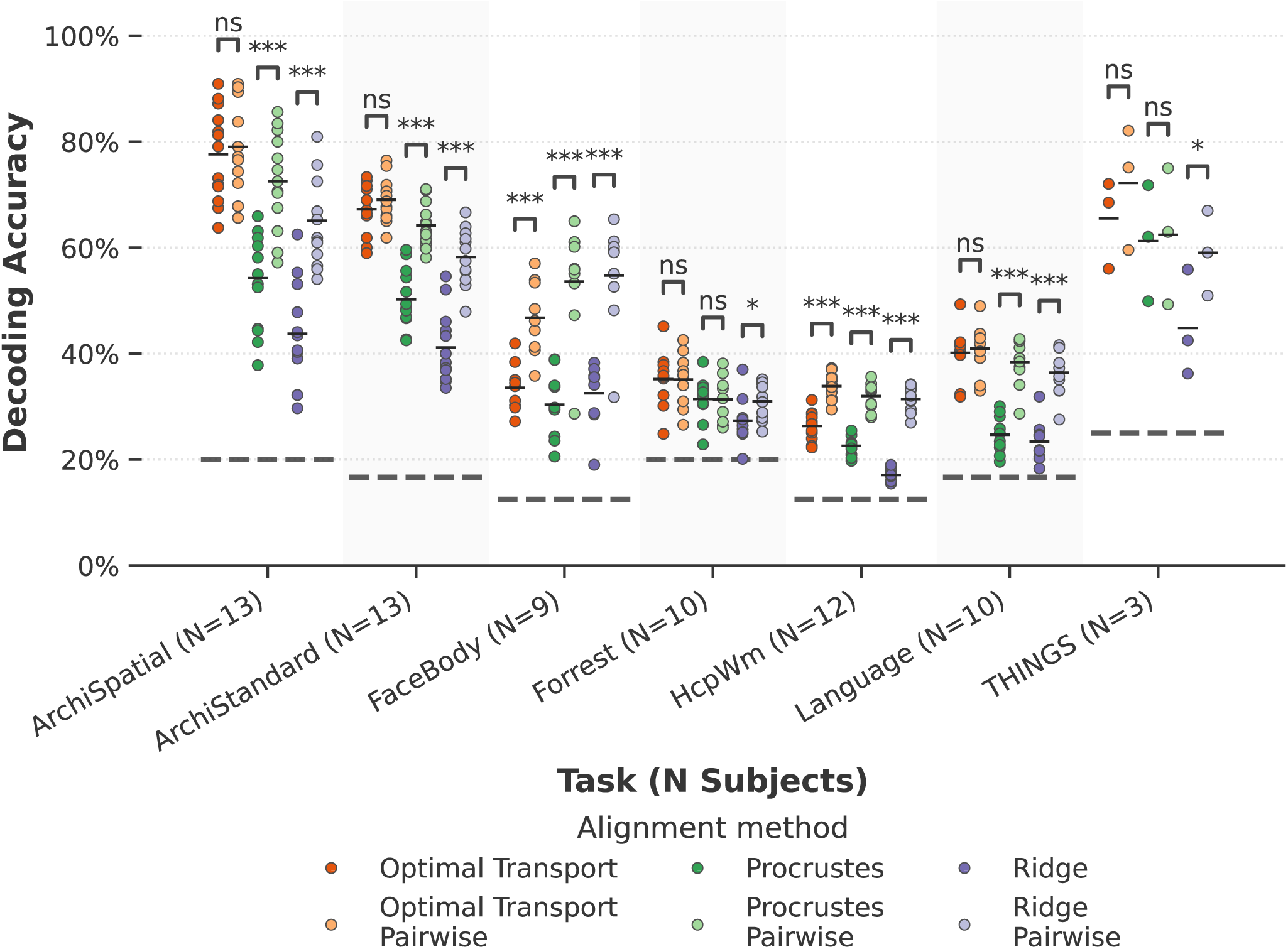
Comparison between Pairwise and Template alignments. Decoding accuracies are compared between two alignment strategies using the same functional alignment algorithms (Optimal Transport, Procrustes and Ridge). In the *(out-of-sample) Template* strategy, templates are generated by leaving aside the test subject. The latter is then aligned in a pairwise fashion to the template. In the *Pairwise* strategy, each subject is sequentially aligned to every other subject individually, and predictions are made on the target subject. Each data point represents the decoding accuracy for a single subject, with colors distinguishing between alignment methods and strategies. Horizontal lines within each distribution represent mean accuracy values. Dashed horizontal lines indicate chance-level performance for each task. Statistical comparisons between in-sample and out-of-sample strategies are performed separately for each alignment method using *t*-tests with Nadeau and Bengio, 1999 correction. Significance levels: *\* p* ≤ 0.05, *\*\* p* ≤ 0.01, *\*\*\* p* ≤ 0.001.

We provide in the appendix the overall computation time for each strategy on Figure 9. We notably observe that Pairwise alignment strategies require less computation time than out-of-sample templates. However, a clear performance gap appears with in-sample-templates that require drastically less computations than out-of-sample templates and pairwise strategies.

### 3.3 OT Templates Show the Smallest In- vs. Out-of-sample Bias

To assess whether incorporating the test subject during template construction introduces a bias, potentially inflating decoding performance, we compare in-sample and out-of-sample template generation strategies (Figure 4).

**Figure 4:**
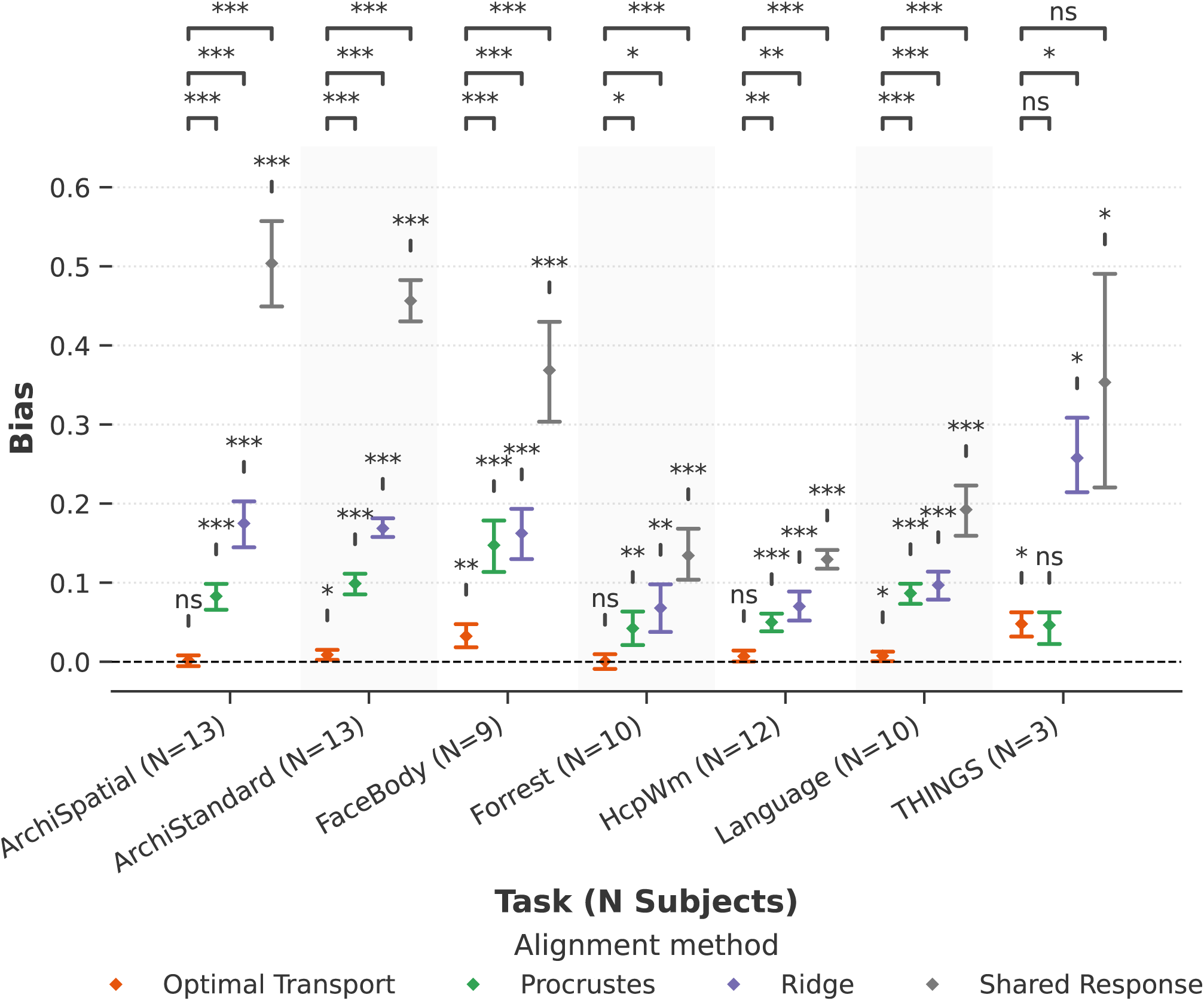
Empirical bias between In-Sample and Out-of-Sample template alignment. Bias is defined as the gap in decoding accuracy between *in-sample* and *out-of-sample* template strategies. The difference is computed for each pair of decoding scores between the strategies for every subject, method and task. In the *in-sample* condition, the template is computed using data from all subjects, including the test subject. In the *out-of-sample* condition, the template is computed excluding the left-out test subject. The test subject is then aligned in a pairwise fashion to the template and predictions are made on it. A one-sample *t*-test with null hypothesis corresponding to the absence of bias (bias = 0) is performed for each method. Results are reported above the error bars. For each task, difference between biases of Optimal Transport and other alignment methods are compared using two sample *t*-tests with Nadeau and Bengio, 1999 correction. Results are reported on the top-row. Significance levels: *\* p* ≤ 0.05, *\*\* p* ≤ 0.01, *\*\*\* p* ≤ 0.001.

This analysis directly parallels the observations of Jeganathan et al., 2024, who reported substantial bias for Procrustes-based templates. In line with their findings, we observe a significant performance increase for Procrustes in the in-sample setting on six of the seven tasks. This confirms that including the test subject in template estimation leads to overly optimistic decoding scores. Ridge behaves similarly: it demonstrates a consistent positive bias across all tasks, with magnitudes exceeding those observed for Procrustes, indicating a greater susceptibility of this method to inclusion bias. SRM follows a similar trend. However, the analysis in Appendix Figure 7 indicates that the large bias mainly arises from the chance-level performance of the out-of-sample template strategy, whereas the in-sample strategy yields decoding scores comparable to those obtained with the other methods. While Chen et al., 2015 reported out-of-subject generalizability of SRM on movie data, our results on task-based data do not provide evidence of such generalizability.

By contrast, Optimal Transport exhibits no significant (*p* ≤ 0.05) bias on three tasks. When a bias is observed, its magnitude is significantly lower (*p* ≤ 0.05) than that obtained with Procrustes and Ridge in every remaining tasks except *THINGS*. Overall, as the number of subjects in a dataset increases, the performance gap between strategies narrows faster for OT than for the other methods.

Taken together, these results suggest that OT-based templates are substantially more robust to the presence of the test subject during template construction. In practical terms, this means that in-sample OT templates can be used with significantly less concern about inflated performance, while also retaining the major computational advantage of avoiding the repeated template estimations required by the out-of-sample strategy.

### 3.4 A Trade-Off Between Spatial Specificity and Activity Variability

With the goal of interpretability in mind, we examine the computed templates and investigate differences in the resulting classifier weight distributions. Since OT, Procrustes and Ridge produce templates in image space, we investigate how they affect decoder feature localization on the cortex.

Since we use linear decoders, we visualize the distribution of voxel contributions in Figure 5, presenting fold-averaged classifier weight maps for the *face-adult* condition of the IBC FaceBody task. Relative to anatomical alignment, Procrustes produces a template where discriminatory voxels (with higher weights in dark red) are clustered on the Fusiform Face Area of the right hemisphere. Ridge regression follows a similar trend, though weights appear more dispersed across the cortical surface.

**Figure 5:**
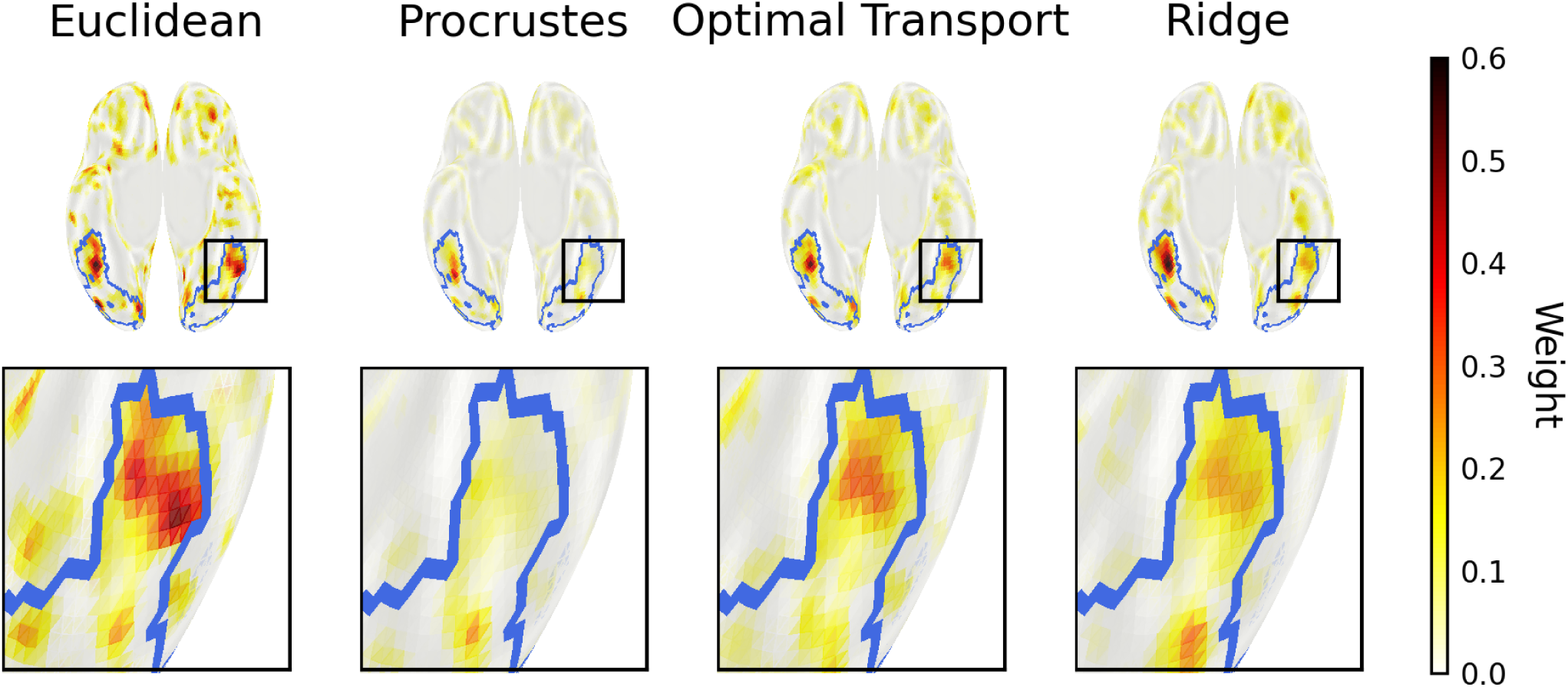
Effect of template alignment methods on decoder weights. We display classifier weights for the *face adult* class against all other conditions from the FaceBody protocol of the IBC dataset as a representative example. Classifier weights are averaged across all alignment folds for each alignment method. The original signals are registered onto the anatomical volumetric MNI152 template and projected onto the *fsaverage5* surface for visualization. **(Top row)**: Medial view of classifier weights on the inflated cortical surface for four alignment methods: anatomical registration only, Procrustes, Optimal Transport, and Ridge. To facilitate comparison, an FDR-corrected cluster analysis (*α* = 0.05) was applied to the group-level statistical map obtained from a fixed-effects analysis across subjects, with results contoured in blue to emphasize the primary region of activity. Black rectangles indicate the zoomed region displayed in the bottom row. **(Bottom row)**: Magnified views of the ventral temporal cortex (black rectangular regions from top row), showing detailed spatial distribution of classifier weights within and around the fusiform region. Warm colors indicate positive weights for the *face adult* class.

In contrast, Optimal Transport shows greater fidelity to the weight distribution of the anatomical template, with clear delineation of discriminatory voxels on the left hemisphere. In addition, the bottom row reveals a stronger spatial specificity with reduced weight spread on the cortical surface compared to the anatomical alignment. Overall, all methods appear to follow the initial topography of the anatomical average, but they differ in how information is distributed across voxels, which underlie the magnitude of the classifier weights.

Moving beyond this specific, qualitative observation, we provide a quantitative analysis of the sparsity across activity maps from different tasks. We compute the ratio between the *L*^2^ and *L*^1^ norm for each aligned map and summarize the values in histograms relative to each method. These results are shown on Fig. 6. Sparse activity maps containing little-to-no activation clusters result in high *L*^2^ to *L*^1^ ratios. We observe that both Procrustes and to a lesser extent Ridge regression exhibit higher ratios, i.e. activity maps are sparser, whereas the distribution under OT template alignment seems to follow more closely the distribution of the anatomical baseline. This implies that OT-projected maps display more structured activity patterns, which corroborates the visual intuition above.

**Figure 6:**
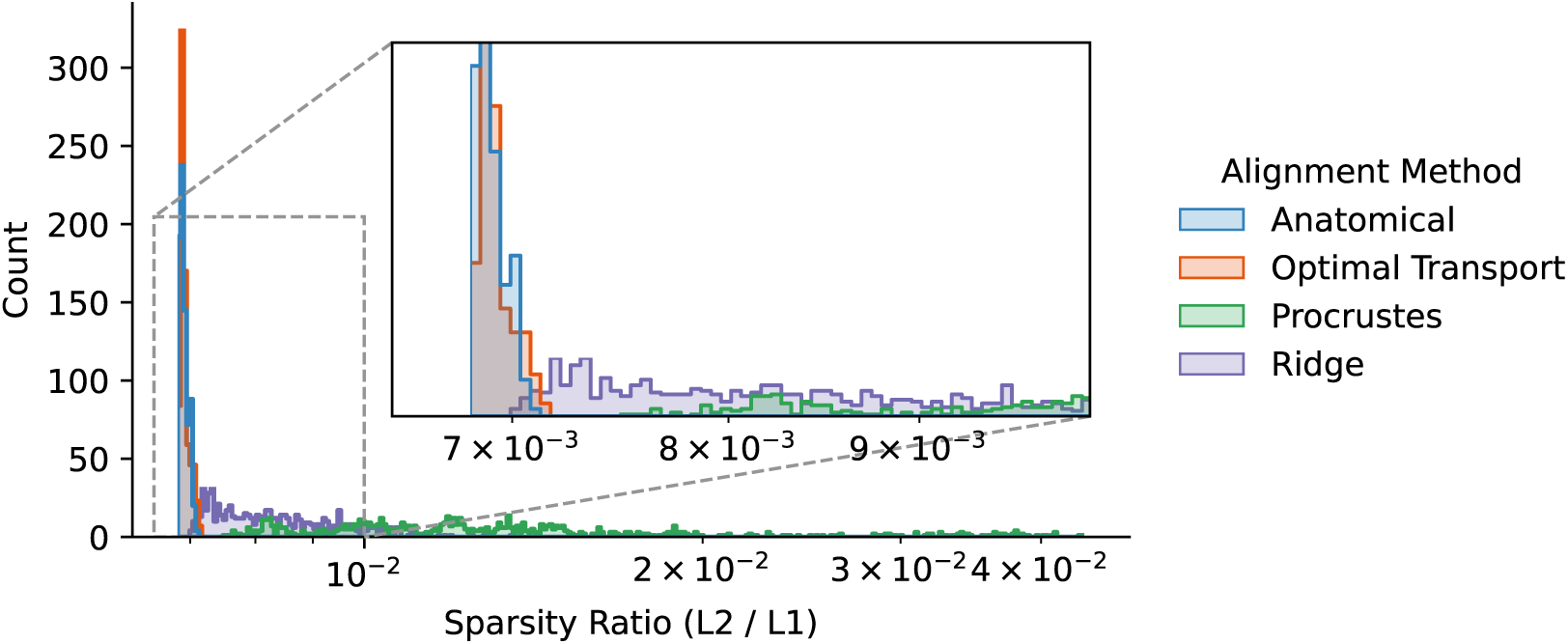
Sparsity profile of activity maps. For each alignment method (anatomical alignment, OT, Procrustes, and Ridge), we compute the ratio of the *L*^2^ to *L*^1^ norm of each projected activity map using the out-of-sample template strategy. We report the distribution of sparsity values for each method as histograms; an inset zooms in on the low-value range to facilitate comparison and axes are set logarithmically. To ensure fair voxel-level comparisons, we restrict the analysis to datasets with 3 mm isotropic resolution, namely the *IBC* and *StudyForrest* datasets.

## 4 Discussion

Let us start by highlighting a few key results of our analysis. We observe a clear performance gap between Optimal Transport and the rest of the methods which often struggle to surpass the anatomical alignment baseline. This gap can be reduced for Procrustes analysis and Ridge regression by using Pairwise or in-sample alignment. However, the Pairwise strategy comes at the penalty of providing no template, and a higher computational cost than the in-sample strategy. Furthermore, the in-sample strategy should be ruled out for Procrustes, Ridge and SRM, given the significant bias related to the inclusion of the test subject when generating the template. The much smaller bias makes the in-sample strategy acceptable for OT. Our examination of the computed templates revealed that while OT, Procrustes and Ridge follow the topography of the anatomical baseline, they exhibit contrasted activity profile, with OT better preserving the anatomical signal distribution, while Procrustes and Ridge tend to nullify larger portions of the activity on the cortex.

### 4.1 A Cross-Validated Framework

In this work, we focus on functional alignment strategies for task decoding. Our alignment/decoding framework is designed to minimize potential sources of variability beyond inter-subject differences. Other benchmarks have relied primarily on movie data (Chen et al., 2015; Guntupalli et al., 2018; Jeganathan et al., 2024) or external tasks (Bazeille et al., 2021), which raises two concerns.

First, discrepancies between alignment and decoding data arise from differences in stimulus type (e.g., movies, resting-state, tasks) and acquisition sessions. This makes it hard to disentangle the effects of inter-subject alignment from other sources of variability on decoding performance.

Second, these approaches have reduced practical applicability, as they require neuroscientists to collect additional data specifically for alignment, which is expensive. Given the cost of fMRI data collection, our 5-fold run-stratified approach makes functional alignment more accessible, as it operates on existing data without additional acquisition costs, assuming that the main task is large enough to support cross-validation splits (which is typically the case in studies designed for decoding analysis).

We also gathered a diverse set of tasks differing in nature, size, and origin. Although hyperalignment has been successfully applied to various movie-viewing datasets (Andreella & Finos, 2022; Haxby et al., 2020), running alignment and decoding in naturalistic setting is relevant only when the amount of data is large (Thual et al., 2023). In our framework, we use a collection of various tasks and datasets with limited amount of subjects and target a comprehensive evaluation of all possible alignment methods and strategies.

### 4.2 Optimal Transport Shows Superior Alignment Performance

Our findings indicate that Optimal Transport alignment achieves the highest decoding performance across all evaluated strategies. Furthermore, at the individual-subject level, OT consistently improves decoding accuracy relative to the anatomical baseline. In contrast, other methods often perform worse than the baseline, thereby defeating the initial purpose of functional alignment.

Our experiments also reveal a minimal gap between in-sample and out-of-sample performance for OT, suggesting little-to-no bias when computing a template at the population level, especially as the number of subjects becomes larger. Specifically, this demonstrates the robustness of the OT templates to the inclusion of a single new subject, and it empirically validates the use of the faster in-sample template strategy with OT, whereas other methods suffer from inflated results due to the inclusion bias.

The possibility of using the in-sample template strategy also facilitates analysis since it yields a unique common space shared by all subjects, making the application of group-level statistical frameworks straightforward. In particular, our analysis of the obtained templates shows that OT is closer to the anatomical template than other methods.

The robust and conservative nature of the OT template stems from the stringent constraints of the transport plans (such as mass preservation), which results in strong sparsity, i.e, only a small number of voxels contribute to the prediction of another one, and in positive coefficients in the alignment matrices (because transport plans encode probability couplings). In contrast to Procrustes rotations or Ridge regression, alignments derived from transport plans preserve small clusters of activity, maintaining the signal’s spatial specificity which drives downstream decoding performance.

Finally, the computational efficiency of the in-sample template OT strategy should be underlined, as it is consistently faster than its pairwise and out-of-sample counterparts, making it highly relevant in practice.

### 4.3 Limitations

Although we sought to encompass several methods and strategies while systematically cross-validating our results to ensure robustness and generalization, several limitations constrain our framework.

We use the Schaefer atlas (Schaefer et al., 2017), following Jeganathan et al., 2024. Control analyses indicate that the number of parcels does not influence our results. Future work could explore alternative atlases or, as suggested by Bazeille et al., 2021, leverage alignment-derived parcellations rather than predefined atlases. Surface-based decoding (rather than the volume-based approach used here) could further improve the correspondence between subjects and reduce mismatches. Our framework, *fmralign*, does not currently support searchlight hyperalignment as in Guntupalli et al., 2016.

Additionally, our approach could be extended to whole-brain alignment techniques that do not rely on local parcels, such as the Promises model Andreella and Finos, 2022 or Fused Unbalanced Gromov-Wasserstein Thual et al., 2022, which incorporate anatomical constraints. However, the choice and tuning of spatial priors in the former framework remain unclear in comparison to parcellated methods, and the latter is computationally prohibitive in the template setting.

Finally, the implementation of pairwise approaches could be accelerated by caching invertible transformations, achieving complexity of *S*(*S* − 1)/2 in the number of subjects. However, we did not implement this strategy because it is not general (e.g. Ridge transformations are not invertible). This optimization could substantially benefit OT and Procrustes. Furthermore, we relied on a CPU implementation of the Sinkhorn algorithm for OT; performance could be dramatically improved using GPU acceleration (Cuturi, 2013).

### 4.4 Conclusion

We develop a comprehensive evaluation of brain alignment techniques demonstrating that in-sample template-based functional alignment coupled with Optimal Transport offers clear improvements in decoding performance compared with other competing methods. Moreover, this strategy is computationally efficient and benefits from stability and parsimony properties that preserve signal specificity, establishing it as a powerful and straightforward tool for neuroimaging research. We carry out our experiments in an open and reproducible fashion, with all methods and strategies available in our package fmralign^2^. All datasets used in this study are publicly available online.

## 5 Data and Code Availability

All functional alignment methods are implemented in the fmralign package, available at https://fmralign.github.io/fmralign/. The code used to reproduce our experiments is publicly available at https://github.com/pbarbarant/fmri template benchmark and is built on the benchopt framework (Moreau et al., 2022). All datasets used in this study are publicly accessible:

- IBC: https://individual-brain-charting.github.io/docs/
- THINGS: https://github.com/courtois-neuromod/cneuromod-things
- StudyForrest: https://www.studyforrest.org/access.html

## 6 Author Contributions

**PL.B**: Conceptualization, Methodology, Software, Formal analysis, Visualization, Original draft, Review and Editing. **F.M**: Conceptualization, Methodology, Review and Editing. **B.T**: Conceptualization, Methodology, Review and Editing.

## 7 Ethics

All datasets used in this study are publicly available and were originally collected as part of previously approved studies conducted independently of the present work.

## A Appendix

**Table 3:** Overview of the selected task protocols across datasets.

| Dataset | Task label | Nature | Subjects | Conditions | Maps/Cond. | Description |
| --- | --- | --- | --- | --- | --- | --- |
| Pinho et al., 2024 | ArchiSpatial | Localizer | 13 | 5 | 16 | Basic cognitive and motor functions using brief visual and auditory stimuli for actions, language, and arithmetic. |
| Pinho et al., 2024 | ArchiStandard | Localizer | 13 | 6 | 20 | Spatial and motor processing through eye movements, grasping, orientation and hand-judgment tasks. |
| Pinho et al., 2024 | FaceBody | Visual | 9 | 8 | 24 | Subjects view images of faces and bodies to examine visual recognition processes. |
| Hanke et al., 2015 | Forrest | Auditory | 10 | 5 | 35 | Subjects listen to the soundtrack of the *Forrest Gump* movie. |
| Pinho et al., 2024 | Language | Reading | 10 | 6 | 120 | Rapid serial visual presentation of words to investigate reading comprehension. |
| Pinho et al., 2024 | HcpWm | Working Memory | 12 | 8 | 3 | Tasks designed to assess short-term memory and cognitive load. |
| St-Laurent et al., 2025 | THINGS | Visual | 3 | 4 | 53 | Subjects are presented with images from the THINGS (Hebart et al., 2019) dataset. |

**Figure 7:**
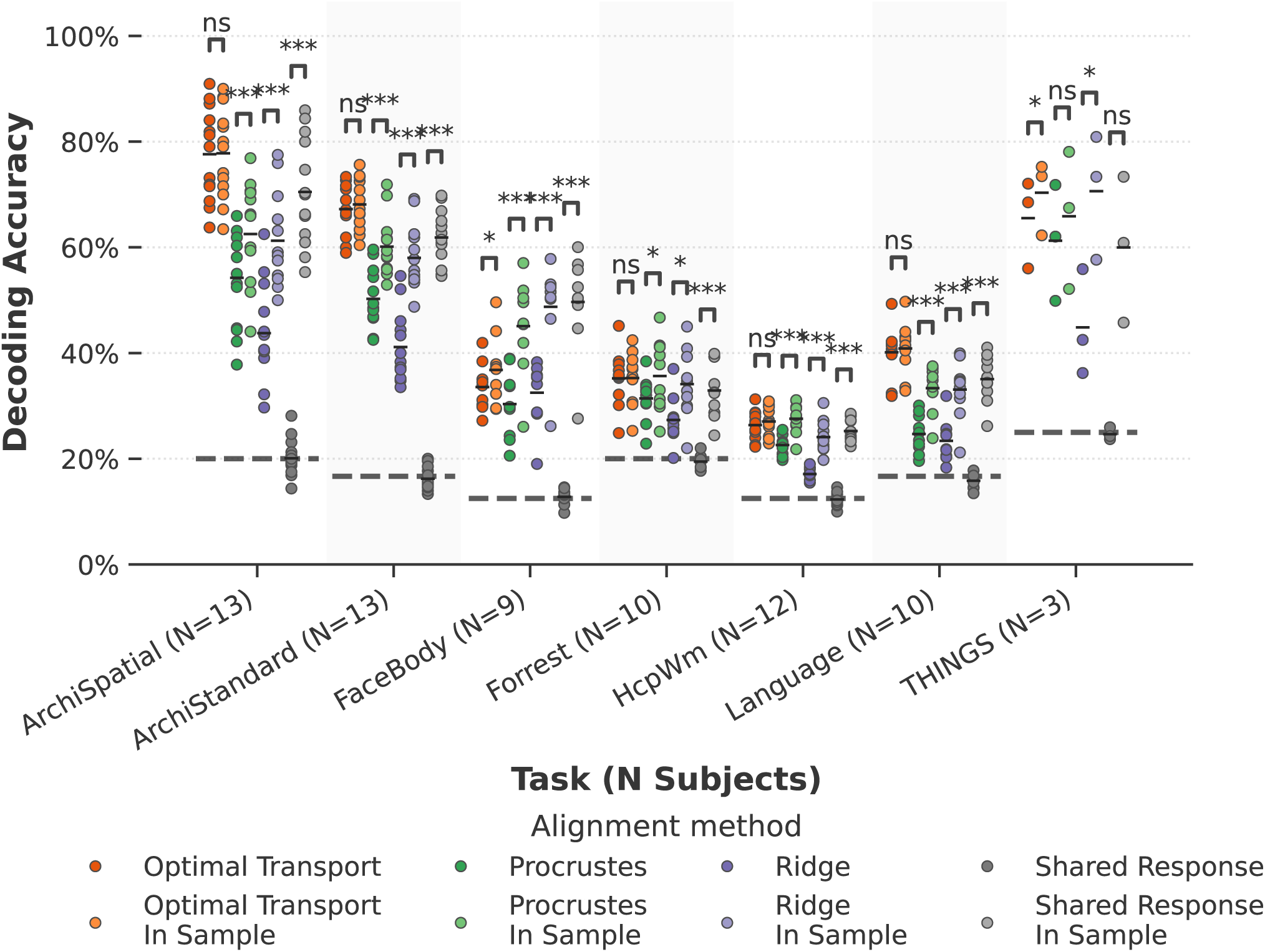
Comparison between In-Sample and Out-of-Sample template alignment. Decoding accuracies are compared between in-sample and out-of-sample template generation strategies, which are fast and slow to compute, respectively. Each data point represents the decoding accuracy for a single subject, with colors distinguishing between alignment methods and template generation strategies. Horizontal lines within each distribution represent mean accuracy values. Dashed horizontal lines indicate chance-level performance for each task. Statistical comparisons between in-sample and out-of-sample strategies are performed separately for each alignment method using *t*-tests with Nadeau and Bengio, 1999 correction. Significance levels: *\* p* ≤ 0.05, *\*\* p* ≤ 0.01, *\*\*\* p* ≤ 0.001.

**Figure 8:**
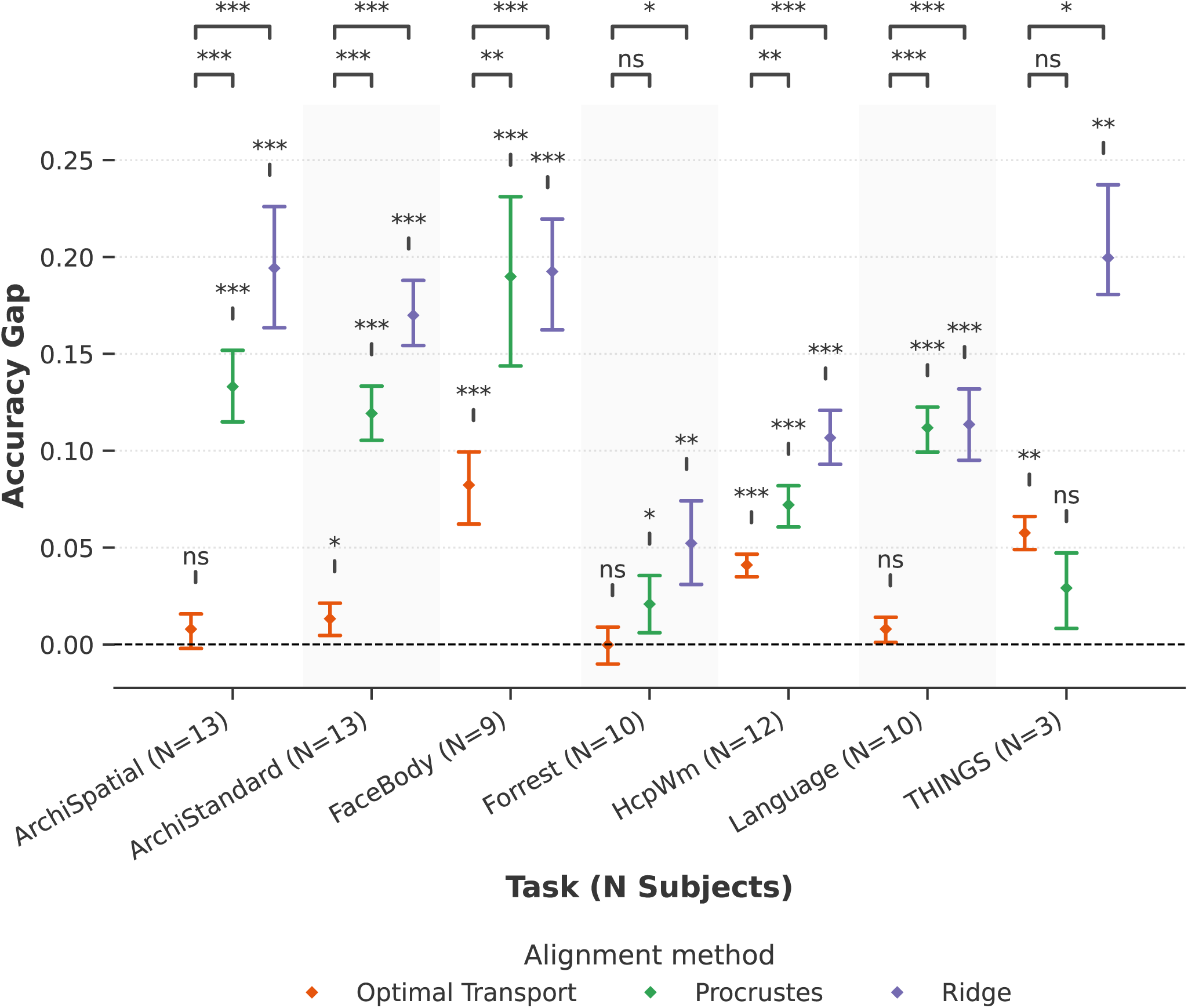
Performance gap between Pairwise and Out-of-Sample template alignment. Accuracy differences between *Pairwise* and *Out-of-Sample* alignment strategies are compared across methods. In the *Pairwise* condition, each held-out subject serves as the alignment target. In the *Out-of-Sample* condition, the template is constructed excluding the test subject, which is then aligned to it in a pairwise fashion before prediction. For each method, a one-sample *t*-test assesses whether the accuracy difference is significantly different from zero; results are reported above the error bars. For each task, difference between biases of Optimal Transport and other alignment methods are compared using two sample *t*-tests with Nadeau and Bengio, 1999 correction. Results are reported on the top-row. Significance levels: *\* p* ≤ 0.05, *\*\* p* ≤ 0.01, *\*\*\* p* ≤ 0.001.

**Figure 9:**
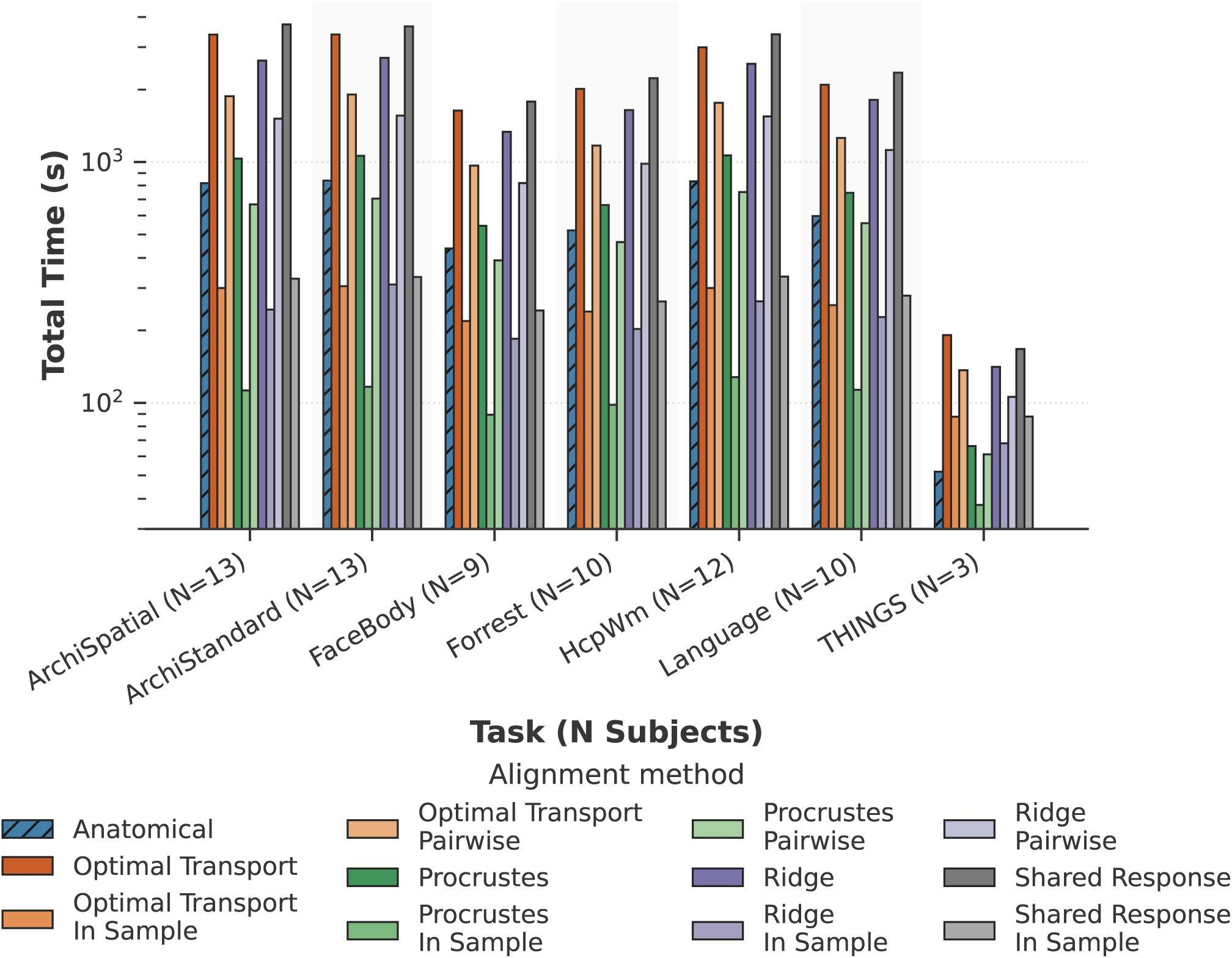
Computation time comparison across alignment methods. Total computation time is reported for each alignment method across different tasks and alignment strategies. Times are summed across all leave-one-subject-out cross-validation folds to reflect the total computational burden for each method-task combination. Bar heights represent the cumulative time (in seconds, log scale) required to align all subjects. Anatomical alignment (hatched bars) serves as a baseline, representing only the time for data processing without functional alignment in the out-of-sample scheme. The default out-of-sample strategy recomputes a template for every test-subject, while the in-sample template-based alignment computes a single population template from all subjects. Pairwise alignment performs individual subject-to-subject alignments for each pair.

**Figure 10:**
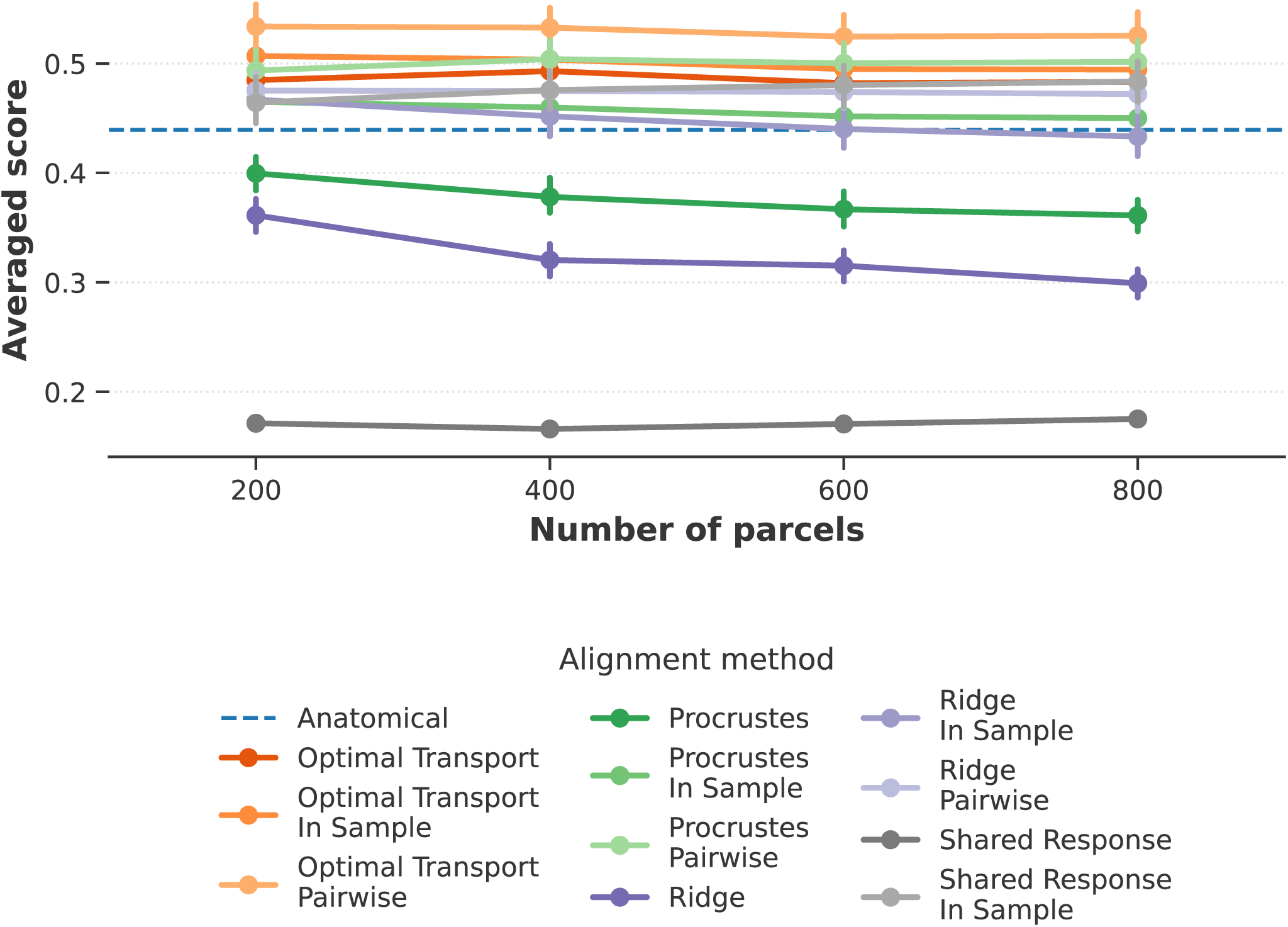
Influence of the number of parcels. Dataset-averaged cross-validated decoding accuracies are shown as a function of the number of parcels, for each alignment method and target strategy. Points indicate mean performance across subjects and folds. Error bars represent the standard error of the mean.

## Footnotes

1 http://fmralign.github.io/fmralign/

2 http://fmralign.github.io/fmralign

## References

Abraham, A., Pedregosa, F., Eickenberg, M., Gervais, P., Muller, A., Kossaifi, J., Gramfort, A., Thirion, B., & Varoquaux, G. (2013). Machine Learning for Neuroimaging with Scikit-Learn. Frontiers in Neuroscience, 15. 10.3389/fninf.2014.00014

Agueh, M., & Carlier, G. (2011). Barycenters in the wasserstein space. SIAM Journal on Mathematical Analysis, 43 (2), 904–924. 10.1137/100805741

Andreella, A., & Finos, L. (2022). Procrustes analysis for high-dimensional data. Psychometrika, 87 (4), 1422–1438. 10.1007/s11336-022-09859-5

Barbosa, J., Nejatbakhsh, A., Duong, L., Harvey, S. E., Brincat, S. L., Siegel, M., Miller, E. K., & Williams, A. H. (2025). Quantifying differences in neural population activity with shape metrics. 10.1101/2025.01.10.632411

Bazeille, T., Richard, H., Janati, H., & Thirion, B. (2019). Local optimal transport for functional brain template estimation. In Lecture notes in computer science (pp. 237–248). Springer International Publishing. 10.1007/978-3-030-20351-118

Bazeille, T., DuPre, E., Richard, H., Poline, J.-B., & Thirion, B. (2021). An empirical evaluation of functional alignment using inter-subject decoding. NeuroImage, 245, 118683. 10.1016/j.neuroimage.2021.118683

Busch, E. L., Slipski, L., Feilong, M., Guntupalli, J. S., Castello, M. V. d. O., Huckins, J. F., Nastase, S. A., Gobbini, M. I., Wager, T. D., & Haxby, J. V. (2021). Hybrid hyperalignment: A single high-dimensional model of shared information embedded in cortical patterns of response and functional connectivity. NeuroImage, 233, 117975. 10.1016/j.neuroimage.2021.117975

Chen, P.-H., Chen, J., Yeshurun, Y., Hasson, U., Haxby, J., & Ramadge, P. J. (2015). A reduced-dimension fmri shared response model. In C. Cortes, N. Lawrence, D. Lee, M. Sugiyama, & R. Garnett (Eds.), Advances in neural information processing systems (Vol. 28). Curran Associates, Inc.

Cuturi, M. (2013). Sinkhorn distances: Lightspeed computation of optimal transport. In C. Burges, L. Bottou, M. Welling, Z. Ghahramani, & K. Weinberger (Eds.), Advances in neural information processing systems (Vol. 26). Curran Associates, Inc.

Ferrante, M., Boccato, T., Ozcelik, F., VanRullen, R., & Toschi, N. (2024). Through their eyes: Multi-subject brain decoding with simple alignment techniques. Imaging Neuroscience, 2. 10.1162/imaga00170

Fonov, V., Evans, A., McKinstry, R., Almli, C., & Collins, D. (2009). Unbiased nonlinear average age-appropriate brain templates from birth to adulthood. NeuroImage, 47, S102. 10.1016/s1053-8119(09)70884-5

Glasser, M. F., Coalson, T. S., Robinson, E. C., Hacker, C. D., Harwell, J., Yacoub, E., Ugurbil, K., Andersson, J., Beckmann, C. F., Jenkinson, M., Smith, S. M., & Van Essen, D. C. (2016). A multi-modal parcellation of human cerebral cortex. Nature, 536 (7615), 171–178. 10.1038/nature18933

Gower, J. C. (1975). Generalized procrustes analysis. Psychometrika, 40 (1), 33–51. 10.1007/bf02291478

Guntupalli, J. S., Feilong, M., & Haxby, J. V. (2018). A computational model of shared fine-scale structure in the human connectome (T. Yarkoni, Ed.). PLOS Computational Biology, 14 (4), e1006120. 10.1371/journal.pcbi.1006120

Guntupalli, J. S., Hanke, M., Halchenko, Y. O., Connolly, A. C., Ramadge, P. J., & Haxby, J. V. (2016). A model of representational spaces in human cortex. Cerebral Cortex, 26 (6), 2919–2934. 10.1093/cercor/bhw068

Hanke, M., Dinga, R., Hausler, C., Guntupalli, J. S., Casey, M., Kaule, F. R., & Stadler, J. (2015). High-resolution 7-tesla fmri data on the perception of musical genres – an extension to the studyforrest dataset. F1000Research, 4, 174. 10.12688/f1000research.6679.1

Haxby, J. V., Gobbini, M. I., Furey, M. L., Ishai, A., Schouten, J. L., & Pietrini, P. (2001). Distributed and overlapping representations of faces and objects in ventral temporal cortex. Science, 293 (5539), 2425–2430. 10.1126/science.1063736

Haxby, J. V., Guntupalli, J. S., Connolly, A. C., Halchenko, Y. O., Conroy, B. R., Gobbini, M. I., Hanke, M., & Ramadge, P. J. (2011). A common, high-dimensional model of the representational space in human ventral temporal cortex. Neuron, 72 (2), 404–416. 10.1016/j.neuron.2011.08.026

Haxby, J. V., Guntupalli, J. S., Nastase, S. A., & Feilong, M. (2020). Hyperalignment: Modeling shared information encoded in idiosyncratic cortical topographies. eLife, 9. 10.7554/elife.56601

Hebart, M. N., Dickter, A. H., Kidder, A., Kwok, W. Y., Corriveau, A., Van Wicklin, C., & Baker, C. I. (2019). Things: A database of 1, 854 object concepts and more than 26, 000 naturalistic object images (F. A. Soto, Ed.). PLOS ONE, 14 (10), e0223792. 10.1371/journal.pone.0223792

Hurley, J. R., & Cattell, R. B. (1962). The procrustes program: Producing direct rotation to test a hypothesized factor structure. Behavioral Science, 7 (2), 258–262. 10.1002/bs.3830070216

Jeganathan, J., Paton, B., Koussis, N., & Breakspear, M. (2024). Integrating anatomical and functional landmarks for interparticipant alignment of imaging data. Imaging Neuroscience, 2, 1–16. 10.1162/imag_a_00253

Kriegeskorte, N., Goebel, R., & Bandettini, P. (2006). Information-based functional brain mapping. Proceedings of the National Academy of Sciences, 103 (10), 3863–3868. 10.1073/pnas.0600244103

Lee, S., Halder, S., Kubler, A., Birbaumer, N., & Sitaram, R. (2010). Effective functional mapping of fmri data with support-vector machines. Human Brain Mapping, 31 (10), 1502–1511. 10.1002/hbm.20955

Moreau, T., Massias, M., Gramfort, A., Ablin, P., Bannier, P.-A., Charlier, B., Dagréou, M., Dupré la Tour, T., Durif, G., F. Dantas, C., Klopfenstein, Q., Larsson, J., Lai, E., Lefort, T., Malezieux, B., Moufad, B., T. Nguyen, B., Rakotomamonjy, A., Ramzi, Z., … Vaiter, S. (2022). Benchopt: Reproducible, efficient and collaborative optimization benchmarks. NeurIPS. https://arxiv.org/abs/2206.13424

Nadeau, C., & Bengio, Y. (1999). Inference for the generalization error. In S. Solla, T. Leen, & K. Muller (Eds.), Advances in neural information processing systems (Vol. 12). MIT Press.

Naselaris, T., Kay, K. N., Nishimoto, S., & Gallant, J. L. (2011). Encoding and decoding in fmri. NeuroImage, 56 (2), 400–410. 10.1016/j.neuroimage.2010.07.073

Pedregosa, F., Varoquaux, G., Gramfort, A., Michel, V., Thirion, B., Grisel, O., Blondel, M., Prettenhofer, P., Weiss, R., Dubourg, V., Vanderplas, J., Passos, A., Cournapeau, D., Brucher, M., Perrot, M., & Duchesnay, E. (2011). Scikit-learn: Machine learning in Python. Journal of Machine Learning Research, 12, 2825–2830.

Pinho, A. L., Richard, H., Ponce, A. F., Eickenberg, M., Amadon, A., Dohmatob, E., Denghien, I., Torre, J. J., Shankar, S., Aggarwal, H., Thual, A., Chapalain, T., Ginisty, C., Becuwe-Desmidt, S., Roger, S., Lecomte, Y., Berland, V., Laurier, L., Joly-Testault, V., … Thirion, B. (2024). Individual brain charting dataset extension, third release for movie watching and retinotopy data. Scientific Data, 11 (1). 10.1038/s41597-024-03390-1

Richard, H., & Thirion, B. (2023). Fastsrm: A fast, memory efficient and identifiable implementation of the shared response model. Aperture Neuro, 3. 10.52294/001c.87954

Sabuncu, M. R., Singer, B. D., Conroy, B., Bryan, R. E., Ramadge, P. J., & Haxby, J. V. (2009). Function-based intersubject alignment of human cortical anatomy. Cerebral Cortex, 20 (1), 130–140. 10.1093/cercor/bhp085

Schaefer, A., Kong, R., Gordon, E. M., Laumann, T. O., Zuo, X.-N., Holmes, A. J., Eickhoff, S. B., & Yeo, B. T. T. (2017). Local-global parcellation of the human cerebral cortex from intrinsic functional connectivity mri. Cerebral Cortex, 28 (9), 3095–3114. 10.1093/cercor/bhx179

St-Laurent, M., Pinsard, B., Contier, O., DuPre, E., Seeliger, K., Borghesani, V., Boyle, J. A., Bellec, L., & Hebart, M. N. (2025). Cneuromod-things, a densely-sampled fmri dataset for visual neuroscience. 10.48550/ARXIV.2507.09024

Thual, A., Benchetrit, Y., Geilert, F., Rapin, J., Makarov, I., Banville, H. J., & King, J.-R. (2023). Aligning brain functions boosts the decoding of visual semantics in novel subjects. ArXiv, abs/2312.06467.

Thual, A., Tran, Q. H., Zemskova, T., Courty, N., Flamary, R., Dehaene, S., & Thirion, B. (2022). Aligning individual brains with fused unbalanced gromov wasserstein. In A. H. Oh, A. Agarwal, D. Belgrave, & K. Cho (Eds.), Advances in neural information processing systems.

